# Inferring gene regulatory networks using transcriptional profiles as dynamical attractors

**DOI:** 10.1101/2023.03.03.530929

**Authors:** Ruihao Li, Jordan C. Rozum, Morgan M. Quail, Mohammad N. Qasim, Suzanne S. Sindi, Clarissa J. Nobile, Réka Albert, Aaron D. Hernday

**Affiliations:** Quantitative and Systems Biology Graduate Program, School of Natural Sciences, University of California, Merced, CA, USA; Department of Systems Science and Industrial Engineering, Binghamton University (State University of New York), NY, USA; Department of Applied Mathematics, School of Natural Sciences, University of California, Merced, CA, USA; Department of Molecular Cell Biology, School of Natural Sciences, University of California, Merced, CA, USA; Department of Physics, Pennsylvania State University, PA, USA; Department of Biology, Pennsylvania State University, PA, USA; Health Sciences Research Institute, University of California, Merced, CA, USA

## Abstract

Genetic regulatory networks (GRNs) regulate the flow of genetic information from the genome to expressed messenger RNAs (mRNAs) and thus are critical to controlling the phenotypic characteristics of cells. Numerous methods exist for profiling mRNA transcript levels and identifying protein-DNA binding interactions at the genome-wide scale. These enable researchers to determine the structure and output of transcriptional regulatory networks, but uncovering the complete structure and regulatory logic of GRNs remains a challenge. The field of GRN inference aims to meet this challenge using computational modeling to derive the structure and logic of GRNs from experimental data and to encode this knowledge in Boolean networks, Bayesian networks, ordinary differential equation (ODE) models, or other modeling frameworks. However, most existing models do not incorporate dynamic transcriptional data since it has historically been less widely available in comparison to “static” transcriptional data. We report the development of an evolutionary algorithm-based ODE modeling approach that integrates kinetic transcription data and the theory of attractor matching to infer GRN architecture and regulatory logic. Our method outperformed six leading GRN inference methods, none of which incorporate kinetic transcriptional data, in predicting regulatory connections among TFs when applied to a small-scale engineered synthetic GRN in *Saccharomyces cerevisiae*. Moreover, we demonstrate the potential of our method to predict unknown transcriptional profiles that would be produced upon genetic perturbation of the GRN governing a two-state cellular phenotypic switch in *Candida albicans*. We established an iterative refinement strategy to facilitate candidate selection for experimentation; the experimental results in turn provide validation or improvement for the model. In this way, our GRN inference approach can expedite the development of a sophisticated mathematical model that can accurately describe the structure and dynamics of the *in vivo* GRN.

**Author Summary:** The establishment of distinct transcriptional programs, where specific sets of genes are activated or repressed, is fundamental to all forms of life. Sequence-specific DNA-binding proteins, often referred to as regulatory transcription factors, form interconnected gene regulatory networks (GRNs) which underlie the establishment and maintenance of specific transcriptional programs. Since their discovery, many modeling approaches have sought to understand the structure and regulatory behaviors of these GRNs. The field of GRN inference uses experimental measurements of transcript abundance to predict how regulatory transcription factors interact with their downstream target genes to establish specific transcriptional programs. However, most prior approaches have been limited by the exclusive use of “static” or steady-state measurements. We have developed a unique approach which incorporates dynamic transcriptional data into a sophisticated ordinary differential equation model to infer GRN structures that give rise to distinct transcriptional programs. Our model not only outperforms six other leading models, it also is capable of accurately predicting how changes in GRN structure will impact the resulting transcriptional programs. These unique attributes of our model, combined with “real world” experimental validation of our model predictions, represent a significant advance in the field of gene regulatory network inference.

## Introduction

Gene regulatory networks (GRNs) are comprised of interactions between sequence-specific DNA-binding proteins, or transcription factors (TFs), and their respective regulatory target genes [1]. The characteristics of GRN stability and adaptability underlie the ability of cells to maintain homeostasis [2], respond to environmental variables [3], develop into multicellular organisms [4], and make cell fate decisions [5]. Inferring the architecture of GRNs based on experimental datasets, also known as the “inverse problem” [6], is important to understanding these cellular processes (see [7, 8] for examples). The advent of high-throughput “omics” techniques [9] has dramatically accelerated the pace by which researchers can obtain these experimental data sets for GRN reverse engineering [10]. A commonly used high-throughput technique is RNA sequencing, which effectively identifies and counts the number of transcripts present for each RNA species, and thus generates a transcriptional profile of the cell or tissue being assayed. With multiple bulk or single-cell transcriptional profiles measured at different time points, or in different cell types, the genes that are upregulated and downregulated can be determined and be used to further infer the logic of the GRNs that underlie those regulatory changes [11]. Although transcriptional profiles are informative and have been widely used to study biological processes of interest, they do not directly reflect the regulatory status of genes (i.e., whether a gene is activated or repressed) [12], since some mRNAs are highly stable and can accumulate in cells, while others are degraded.

In the past twenty years, many modeling approaches have been developed to infer GRN architectures using “omics” data [9, 13–15]. GRN inference models can be broadly categorized into three distinct categories, based on the algorithms and hypotheses they employ (see reviews: [16–24]): (i) data-driven static models, which do not simulate the biological processes such as transcription or translation, but hypothesize that interacting genes have correlated expression and use the correlations to infer GRN architecture [25, 26]; (ii) discrete models, which simulate the time evolution of discrete variables that qualitatively describe the activity of genes [27, 28]; and (iii) continuous dynamical models which simulate the dynamics of gene expression processes in a quantitative manner based a set of linear [29] or non-linear [30, 31] ordinary differential equations (ODEs).

Dynamical models suffer from the “curse of dimensionality”, i.e., the problem of a state space or parameter space growing exponentially with the number of genes considered. One approach to dealing with this challenge when building dynamical models is to focus on dynamics near attractors [32–34], the states toward which a kinetic system tends to evolve and converge. This approach has been successfully applied in Boolean network inference of GRNs [34–38], in which binary variables, 1 and 0, define the state of a gene as “on” or “off”, respectively. Here, we focus on extending the attractor matching approach from Boolean models to ODE models, which can simulate continuous gene expression levels. Specifically, we infer an ODE model of a GRN whose attractors match the experimentally measured attractor states. The main obstacle to this attractor matching approach to network inference is the lack of experimentally measured GRN kinetic data [23, 39], such as the rates of transcription and mRNA degradation, which shape the dynamics of a GRN system and are essential for determining the attractors’ positions and basins [12]. As a result, most ODE models must rely on fitting procedures to estimate kinetic parameters [40–44]. This strategy differs significantly from the application of attractor dynamics, where measured kinetic parameters are already known and are used to find and match the attractors. Thus, there are few GRN inference techniques available that can effectively leverage kinetic data when it is available. Although the attractor matching approach has not been applied to ODE-based models for the purpose of GRN inference, previous work has explored the potential of attractor-matching strategies in ODEs [45, 46], and several software tools have been developed in this area. For instance, FOS-GRN [47] and Netland [48] can reconstruct multi-attractor kinetic landscapes with ODEs and user-defined parameters. In addition, there are several studies [49–55] exploring how ODE models of GRNs can be steered from one attractor to another; many of these techniques have Boolean analogues, which led us to more closely examine attractor matching inference strategies for ODEs.

In this work, we have developed a GRN inference approach that extends the application of the attractor matching strategy from a Boolean model to an ODE-based model by incorporating measurements of mRNA synthesis and degradation. Our model can simulate genetic regulatory processes with a novel parameterization framework that is built using combinatorial logic operators and Hill functions and can reveal the correlation between a GRN architecture and its dynamics in terms of fixed-point attractors. We have tested our model using both *in silico* data and experimental data generated from a real-life GRN constructed using synthetic biology [56]. Our results show that the use of kinetic parameters and the application of attractor dynamics can significantly improve the inference performance of ODE-based GRN models. Furthermore, since the model simulates GRN dynamic systems in a quantitative manner, it can also predict stable-state transcriptional profiles when given a GRN architecture and kinetic parameters. Using this model, we are able to estimate, for the first time, the similarities between an unknown real-world GRN of an organism and the inferred GRN model based on predictions of the steady state transcriptional profiles that result under perturbations that were not incorporated into the inference process.

## Methods

### GRN model architecture

Drawing on the conventions of early work, we depict the GRN architecture as a directed graph consisting of nodes representing both TF proteins and their respective coding genes, and edges representing interactions among these nodes. Non-TF coding genes are not considered in our model. For example, in a simple three gene GRN architecture (Fig. 1), the nodes A, B, and C represent three distinct TF proteins and the genes that encode them, while the connecting lines indicate physical interactions between the TF protein(s) and their respective regulatory target genes. In our framework, two types of interactions exist between TFs and genes: activating or inhibitory, represented by pointed or blunt arrows, respectively. As shown in Fig. 1b and 1c, we denote these TF-gene relationships by an adjacency matrix (named *AM*), which uses 1 or −1 to indicate the activating or inhibitory interactions, and 0 to indicate the absence of an interaction between a given TF and target gene. In addition to the TF-gene interactions, TF-TF interactions may also exist, and are represented by the logic operators AND, OR, or NOT, which qualitatively indicate how multiple TFs are aggregated to affect a common target gene. The qualitative logic of these TF-TF interactions is organized in a protein coordination matrix, or set of logic gates (denoted by *LG*), whose Boolean values are assigned to decide whether the activators or repressors of a gene work synergistically or independently. If a gene in the model has fewer than two TFs, its protein coordination parameters do not affect the GRN dynamics. In this case, we fix the protein coordination parameters to 0. We use *f*_0_, which is bounded between 0 and 1, to represent the basal expression level of a gene when no TF acts upon it. The overall GRN architecture, or *A_net_*, can be expressed by *AM_n×n_*, with *LG_2×n_* and *f*_0_ as the features of the network kinetics, where *n* denotes the number of genes in the GRN. In this simplified GRN architecture, the activators or repressors of a gene work either synergistically or independently (Fig. 1c). This approach greatly reduces the complexity compared to fully general network logic gates (2*N* binary degrees of freedom per node), but leaves many degrees of freedom (*N*^2^ trinary degrees of freedom) in the GRN topology. For example, a GRN consisting of 3 genes has 3^9^ possible configurations in AM and 2^6^ in LG. The qualitative character of the GRN architecture will systematically be made quantitative, as we describe in subsequent sections.

**Figure 1:**
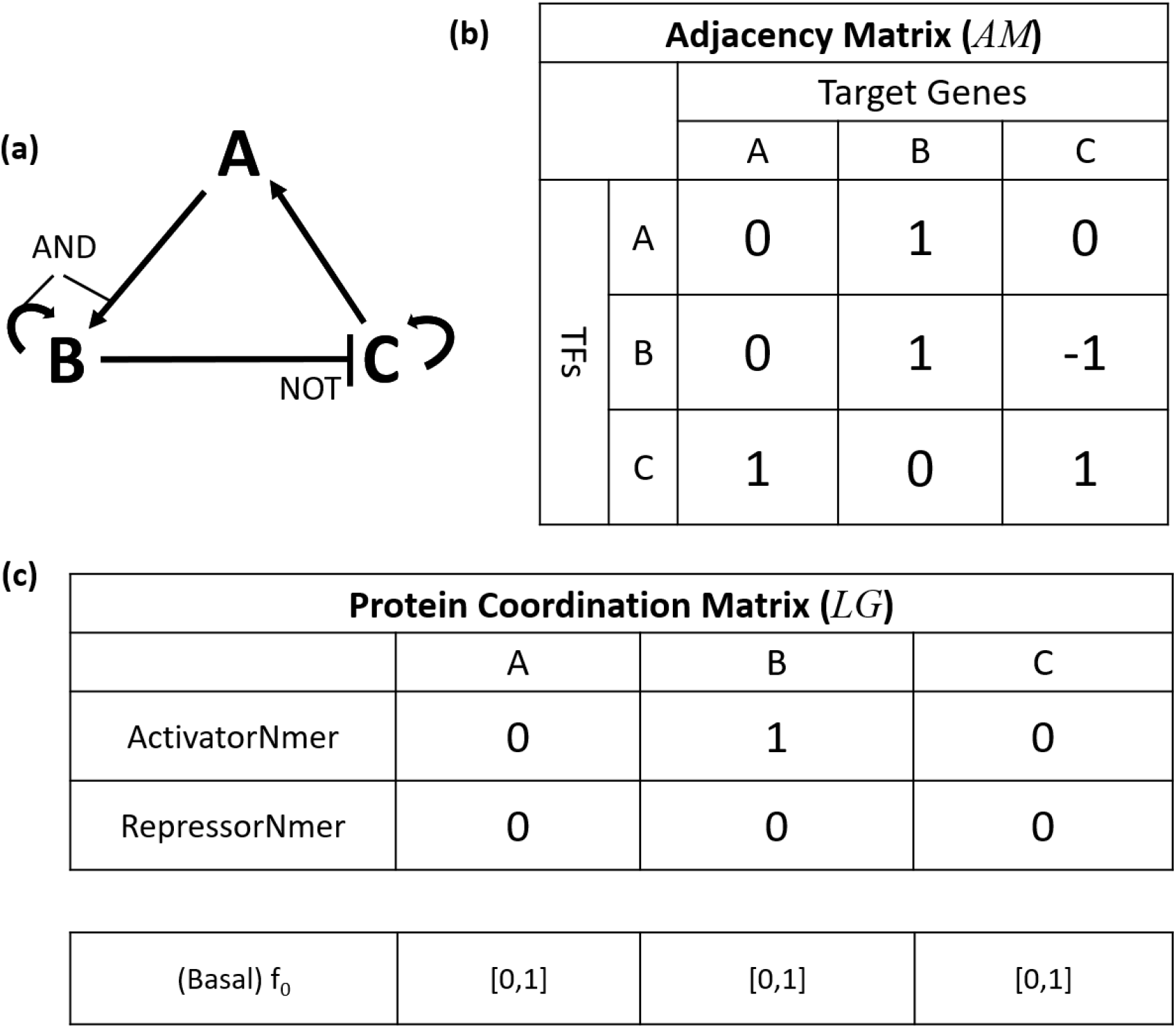
Depiction of a hypothetical GRN architecture. (a) Schematic of a simple GRN in which A and B cooperatively activate B, C activates A and itself, and B represses C in a manner that can override the self-activation of C. (b) The network topology table represents the direct activating, inhibiting, and null connections by 1, −1, and 0, respectively. (c) The protein coordination parameters are assigned to each gene in the genome and qualitatively describe the coordination between each gene’s regulatory TFs. ‘ActivatorNmer’ decides whether the activators of a gene work independently (0) or cooperatively (1); ‘RepressorNmer’ decides whether the repressors of a gene work independently (0) or cooperatively (1). *f*_0_ determines the basal expression level of a gene and whether its activators or repressors outcompete the other.

### Overview of the GRN dynamic system

A list of symbols and parameters used in the GRN dynamic system is given in Table S1. We assume that the diffusion and binding processes of TFs, which happen much faster than transcription and translation, are instantaneous. We assign a *V_max_* and a *V_min_* to each gene, which represent the potential highest and lowest production rates of mRNA transcripts, respectively. We assume linear decay of the proteins and mRNA, and linear translation. These assumptions give rise to Eq. 1 and 2:

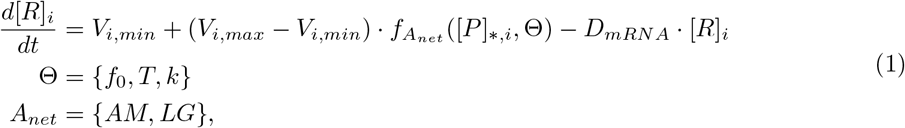

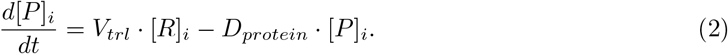

In these equations, [*P*]_*,*i*_ represents the concentrations of the TFs that regulate the *i^th^* gene in the GRN, and *f_A_net__* is the regulation function, which describes how TFs regulate a gene. Other symbols in Eq. 1 and 2 are defined in Table S1. The regulation function *f_A_net__* is a continuous function given in Eq. 9. Typically, Hill functions [57–59] and sigmoid functions [60, 61] form the building blocks of the regulation function. We use Hill functions (Eq. 3 and 4) to formulate the regulation function *f_A_net__*:

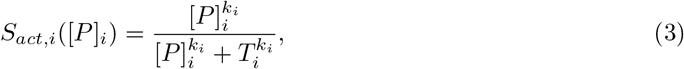

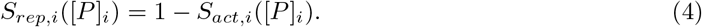

In Eq. 3 and 4, *T* is the protein abundance producing half occupation (disassociation constant of the TF binding) and *k* is the Hill coefficient (effective cooperativity). Eq. 3 describes the regulatory contribution of an activator, and Eq. 4 serves the same role for a repressor. The effect of multiple TFs on the transcription rate, as shown in Fig. 2 a and b, is determined by the choice of *LG* according to the combinatorial Hill functions equations below (Eq. 5 to Eq. 8):

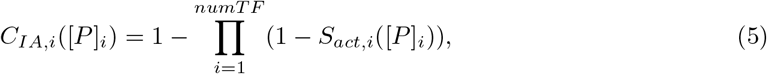

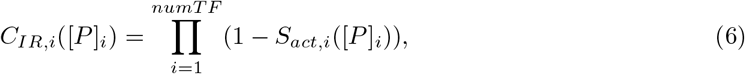

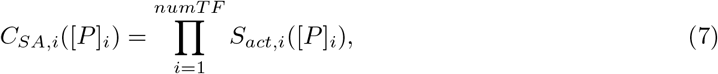

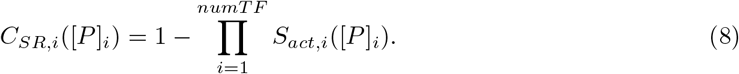

*C_IA,i_*, *C_IR,i_*, *C_SA,i_*, and *C_SR,i_* denote the combinatorial Hill functions for independent activators, independent repressors, synergistic activators, and synergistic repressors, respectively.

**Figure 2:**
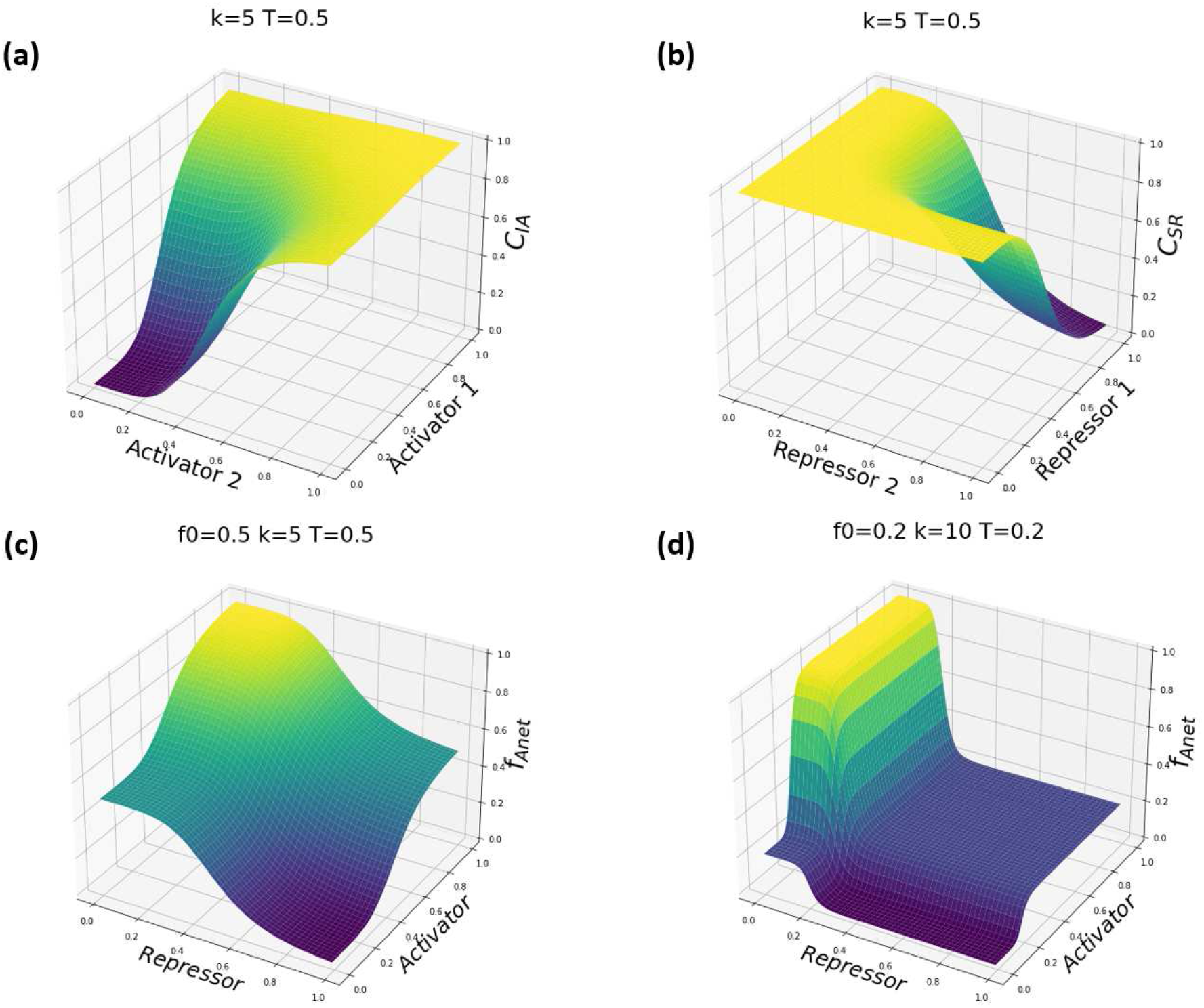
Demonstration of the combinatorial Hill functions (a and b), and the regulation function *f_A_net__* (c and d) under the regulation of an activator and a repressor. In the top two panels (a and b), two combinatorial Hill functions are plotted; in (a) two activators work independently to activate a target gene, while in (b), two repressors work synergistically. In the bottom two panels (c and d), the dependence of the regulatory function *f_A_net__* on the activating and repressing combinatorial Hill functions is plotted for two example cases. In (c), *f_A_net__* achieves the basal transcription rate fraction of 0.5 when there is a lack of both activator and repressor, or when both are present. Activation (resp. inhibition) occurs when the activator (resp. repressor) is abundant, and the repressor (resp. activator) is scarce. In (d), The Hill coefficient, *k*, determines the steepness of the regulation function; The basal expression level, *f*_0_, controls the position of the middle plane and can slide between 0 and 1. The threshold *T* decides the TF abundance that will trigger the activation or repression.

With the *A_net_* and the abundances of the TFs, we define the regulation function *f_A_net__* by Eq. 9, which applies combinatorial Hill function formulas and *f*_0_ to represent the gene regulations:

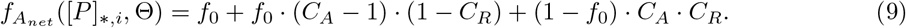

Here, [*P*]_*,*i*_ denotes the effective abundance of the TFs regulating the *i^th^* gene. For notational convenience, we define *C_A,i_* = *C_SA,i_* if activators are synergistic and *C_A,i_* = *C_IA,i_* if they are independent, and similarly for *C_R,i_*. How the parameters affect the shape of *f_A_net__* is shown in Fig. 2 c and d.

### GRN architecture inference

With the deterministic GRN dynamic model constructed above, we propose to infer the *A_net_* using experimentally observed data. Specifically, we assume that the transcriptional profiles of cells in the exponential phase of growth under defined stable culture conditions can be considered as fixed-point attractors in the GRN dynamic system. By incorporating experimentally determined transcription, translation, and degradation rates, we simulate the GRN dynamics and determine whether a given *A_net_* can accurately reproduce the observed attractors. To search for the optimal *A_net_* for a particular GRN, we utilize a modified evolutionary algorithm [62, 63] to iteratively refine the *A_net_* parameters until the predicted network attractors converge upon the experimentally measured ones.

Since *A_net_* and the values of some system parameters are often unknown in practice, we will make use of the measurable kinetic parameters (including *V_max_*, *V_trl_*, *D_mRNA_*, and *D_protein_*) and the steady-state transcriptional profiles to estimate the unknown parameters (*V_min_*, *T*, *k*, and *f*_0_) for a given network architecture (See Table S1). First, the *V_min_* value for a gene is estimated by the minimal expression level of the gene across all the samples. Second, without other prior knowledge, we must assume that when the genes are under TF regulation, they have the same chance to be activated or inhibited. Hence, the *T* of the TFs is calculated by their average expression levels using Eq. 2. Third, we assume that the input transcription profiles include fully activated genes, and estimate *k* by invoking the steady state assumption for transcriptional profiles that include maximal expression. The last unknown parameter, *f*_0_, is obtained by solving Eq. 1, assuming the inputs are at steady state and averaging across inputs. The equations for solving these parameters can be found in the supplemental material (Eq. S1-Eq. S7).

The modified evolutionary algorithm that we developed to efficiently search the GRN architecture space is illustrated in Algorithm 1. First, we randomly create a population of *A_net_*, each of which has the same initial fitness score. Second, we update the estimates of the unmeasured parameters for each *A_net_*. If an *A_net_* has a *f*_0_ less than 0 or greater than 1, a penalty will be added to the fitness score for *A_net_*. Third, we examine each independent self-activating edge in an *A_net_*. If any input with the self-activating gene does not have a companion input state with that gene active, we construct such a state for use as an initial condition to test whether such a state will converge into a measured attractor. When the system is initiated in such a state, the input transcriptional profile suggests that the self-activating gene should eventually be inactivated by other TFs. By including these additional initial conditions, we penalize networks that are highly stable for all or nearly all initial states. Next, we add a mild perturbation to each of the initial states and numerically integrate the deterministic GRN dynamical system using the Runge-Kutta 4^*th*^ order method [64, 65] to find the final steady states. Since we assume that the measured transcriptional profiles represent attractors, these initial states should converge to the corresponding attractor states. If the system state ends up far away from the attractors, the current *A_net_* cannot generate the anticipated attractors. We use a normalized Manhattan distance between the attractors and the final states as a metric:

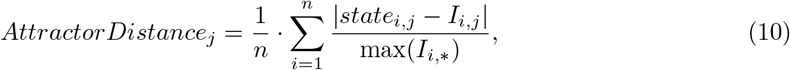

where *state_i,j_* is the *i^th^* gene expression level of the *j^th^* final state obtained by running the GRN dynamic system, *I_i,j_* is the *i^th^* value of the *j^th^* input state, *m* is the total number of attractors, and *n* is the total number of genes.

#### Algorithm 1 Evolutionary Algorithm

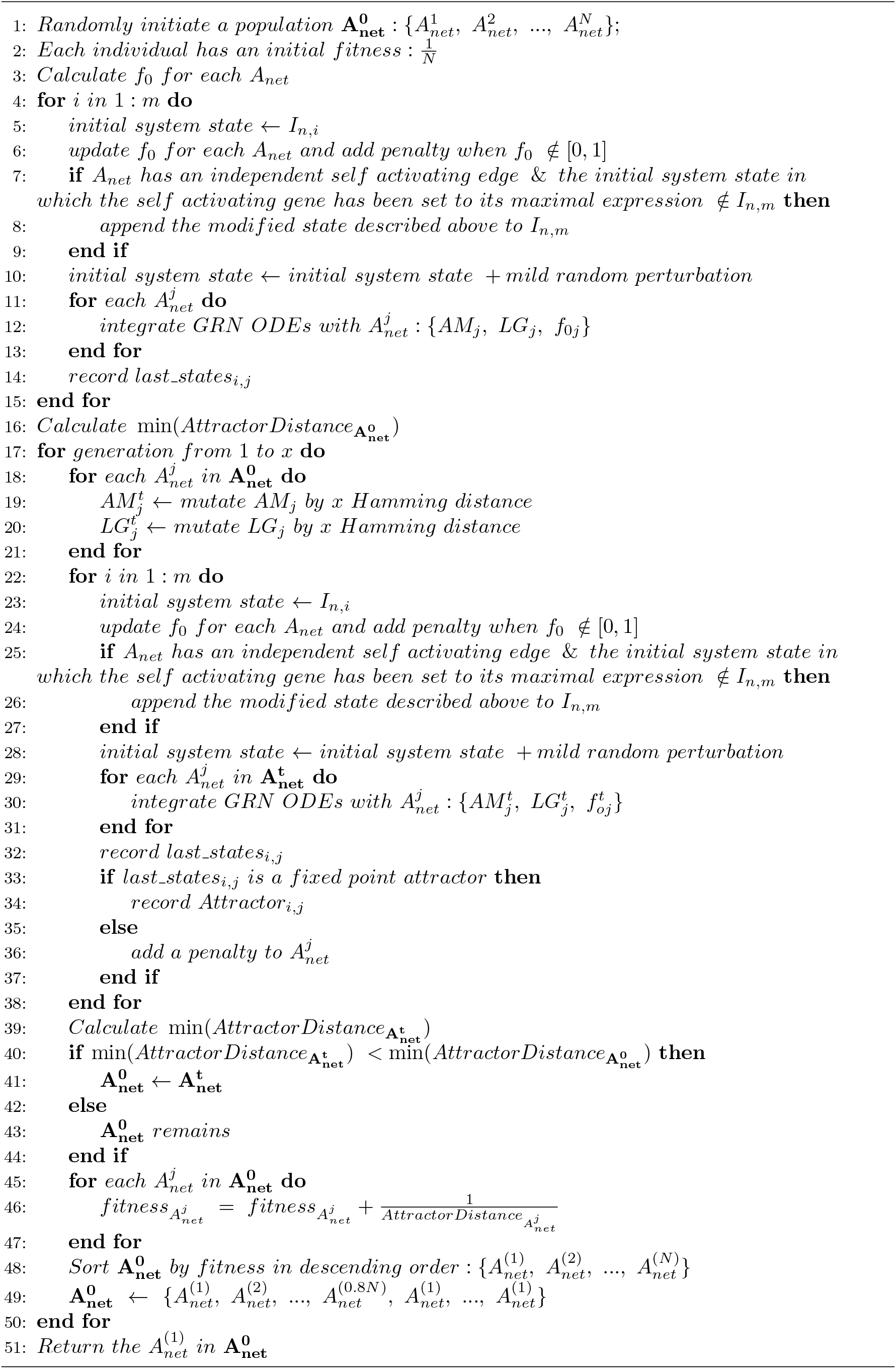

Autoregulation tends to produce excessive attractors not belonging to the input. We compensate for this by considering additional initial states. If one of these additional initial states belongs to the input, the final state will be compared against the initial state. Otherwise, the final state will be compared against the closest attractor in the input. If the initial condition does not converge to a steady state, we apply a penalty to the fitness of the network. We define the fitness of each *A_net_* in the current population by the reciprocal of the minimal average “AttractorDistance” for all initial conditions, subtracting off the penalties for steady state convergence failure and for unrealistic estimation of *f*_0_.

After the attractor distance is calculated, we create the next generation population by randomly mutating each *A_net_* by a certain Hamming distance (see supplemental material), and numerically solve the new ODEs constructed by the mutated *A_net_* to obtain the new minimal average “AttractorDistance”. If the new minimal average “AttractorDistance” is less, we keep the mutated population. Otherwise, the mutated population will be abandoned, and we will return to the former population. Finally, we update the fitness of each *A_net_* according to their “AttractorDistance”. We sort the population by fitness in descending order and then eliminate the last 20% of individuals and duplicate the fittest 20% to restore the population number. By iteratively running the algorithm, we can obtain the fittest *A_net_* in the last generation as the output.

Because the sampling and mutating steps in the algorithm are stochastic, the output *A_net_* can be different each time. We draw a consensus GRN architecture with the assumption that a particular regulatory connection (i.e., an entry in *A_net_*) occurring at a higher frequency among a group of inferred GRN architectures is more significant than one occurring at a lower frequency. In all results presented here, unless otherwise specified, we conducted 30 independent and identical GRN inference processes and obtained a consensus GRN architecture by averaging the fittest *A_net_* architectures, weighting each by its fitness.

In addition to regular transcriptional profiles, our approach can also incorporate data from genetically engineered strains. For instance, if the input transcriptional profiles are from knockout or overexpression strains, we can set the mRNA abundance of the inoperative genes to zero or maximal expression level during the numerical integration. If a regulatory connection is removed by genetic engineering (e.g., by disrupting a TF binding site in a promoter), we can fix the corresponding entry in *A_net_* as zero to eliminate the effect of the disrupted regulatory connection. Furthermore, we can use genome-wide binding data, such as chromatin immunoprecipitation (ChIP) data, to guide the mutation of *A_net_* during inference; if a physical binding interaction exists between a TF and a gene, it is likely to represent a regulatory connection. Therefore, we can lower the probability of the corresponding entry being mutated to zero. The accommodation of different data types allows the model to integrate more available data and perform better.

### Model networks for validation

It is extremely challenging to directly determine the complete and comprehensive composition and structure of “real-world” GRNs in living organisms [18, 66]. Therefore, the use of experimental data in GRN inference can be problematic when it comes to validating the outcome of GRN model predictions, since one can rarely, if ever, be certain that the experimental data provides a complete picture of the real-world GRN structure. For this reason, it has become common practice in the field of GRN inference to utilize *in silico* (i.e., computer generated) datasets for method validation, which can provide gene expression data that is directly predicted based on a hypothetical “source” GRN model [57–59, 67].

We used both *in silico* and biologically observed GRN instances to evaluate our approach. The *in silico* instance consists of five arbitrarily generated toy GRN ODEs (Fig. 3a), each of which has at least 9 different fixed-point attractors. The *A_net_* architectures for these systems are regarded as the reference GRN architectures, which will be used as answer keys against which the inferred GRN architectures will be compared. The *in silico* fixed-point attractors for each *A_net_* are generated by SynTReN, a commonly used benchmark generator for GRN inference [57]; this is done so that the results are not biased by generating state data using the ODE framework we aim to evaluate. Considering that noise generally exists in the experimentally derived transcriptional profiles, we added a Gaussian distributed noise to the *in silico* input transcription profiles. The kinetic parameters of the model were assigned by the values experimentally measured in *Escherichia coli* [68–71] (See Table S2).

**Figure 3:**
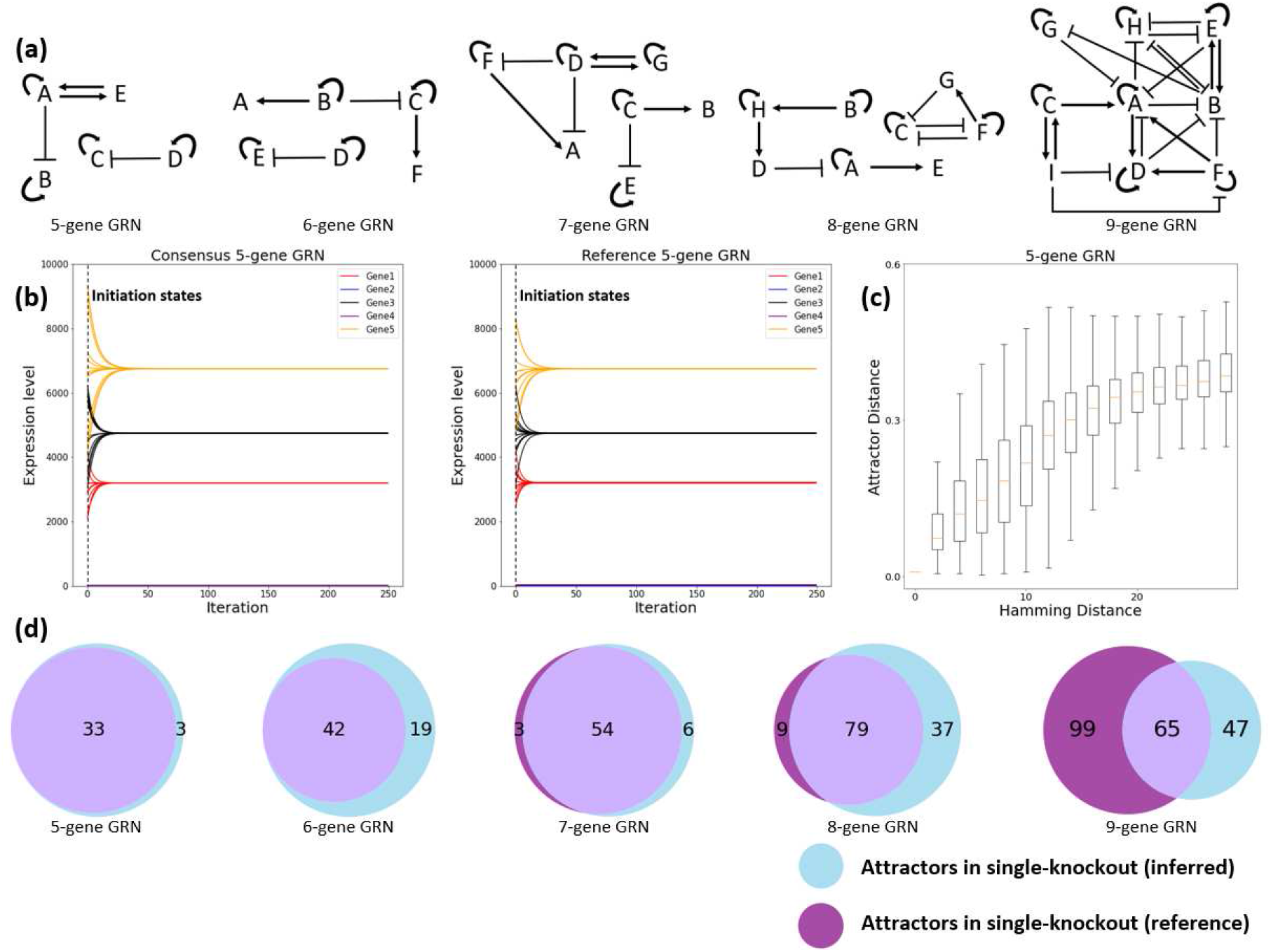
(a) Five GRN architectures were arbitrarily generated as references in the *in silico* test. They have 5-9 genes and at least 9 different fixed-point attractors. The pointed and blunt arrows represent activating and repressing regulatory interactions, respectively. (b) GRN dynamics when initiated near a fixed-point attractor of a reference GRN. The consensus GRN was inferred by the attractors of the 5-gene *in silico* reference GRN. The initial states were obtained by the attractor position plus a Gaussian distributed random variable. The dynamics showed similarly good agreement in other reference GRNs. (c) Positive correlation between *A_net_* similarity (Hamming distance on the horizontal axis) and attractor profiles similarity (attractor distance on the vertical axis). Each column in the box plot contains 1000 random 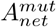 mutated from the 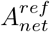 consisting of 5 genes. Similar strong correlation has been observed in all other reference GRNs (see Fig. S1). (d) *in silico* attractors prediction result summary. Two attractors considered matched have an attractor distance less than 0.16 (a cutoff below which a simple null model has a less than 5% chance of producing matched attractors; see Table S4). Overall, the single-knockout reference GRNs produced 384 fixed-point attractors and the single-knockout inferred GRNs produced 385. Of these attractors, 273 were matched. No attractors were matched in a random GRN.

To test our model against an *in vivo* GRN instance, we used experimental data derived from a synthetic GRN engineered in *Saccharomyces cerevisiae* by Cantone et al. [56]. Consisting of 5 genes and a variety of regulatory interactions (Fig. 5 a), the GRN can switch amongst 10 distinct stable states in response to the overexpression of each individual gene in two different carbon sources, galactose and glucose. These stable states were measured by quantitative PCR (qPCR) and converted to absolute expression levels. The promoter strengths, which indicate the rates of transcription initiation for each gene in the GRN, have been estimated by a stochastic optimization algorithm from steady-state gene expression data measured by qPCR [56]. Other kinetic parameters used in the *in vivo* test are provided by Table S2. The *A_net_*, transcription profiles, and kinetic parameters for the *in silico* and *in vivo* tests can be found in the supplemental material.

**Figure 4:**
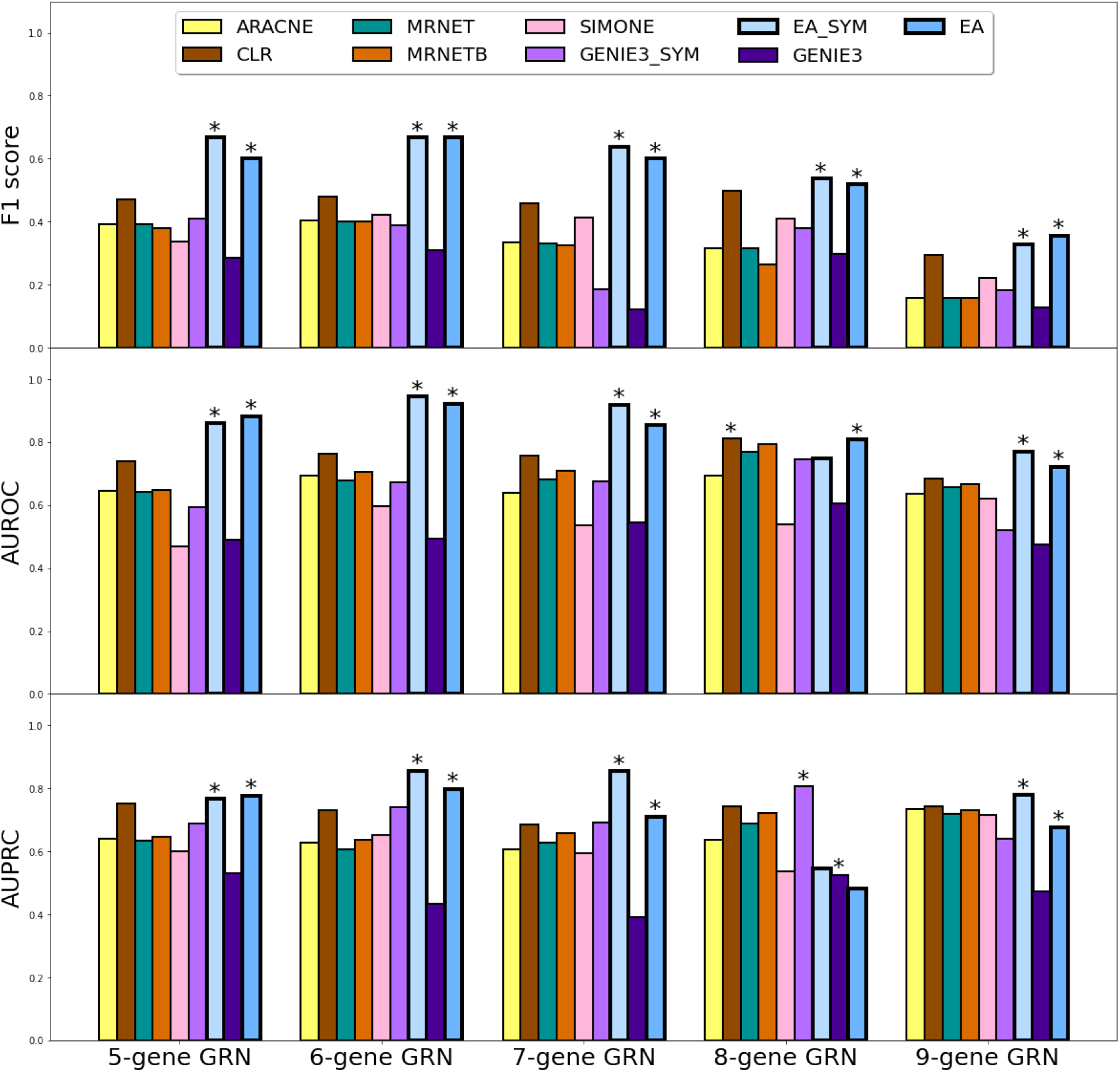
The in *silico* test comparison result in F1 score (upper panel), AUROC (middle panel), and AUPRC (bottom panel). The F1 scores are calculated using a threshold cutoff of 0.5 for all models. Best performances are marked by asterisks for symmetric and asymmetric methods.

**Figure 5:**
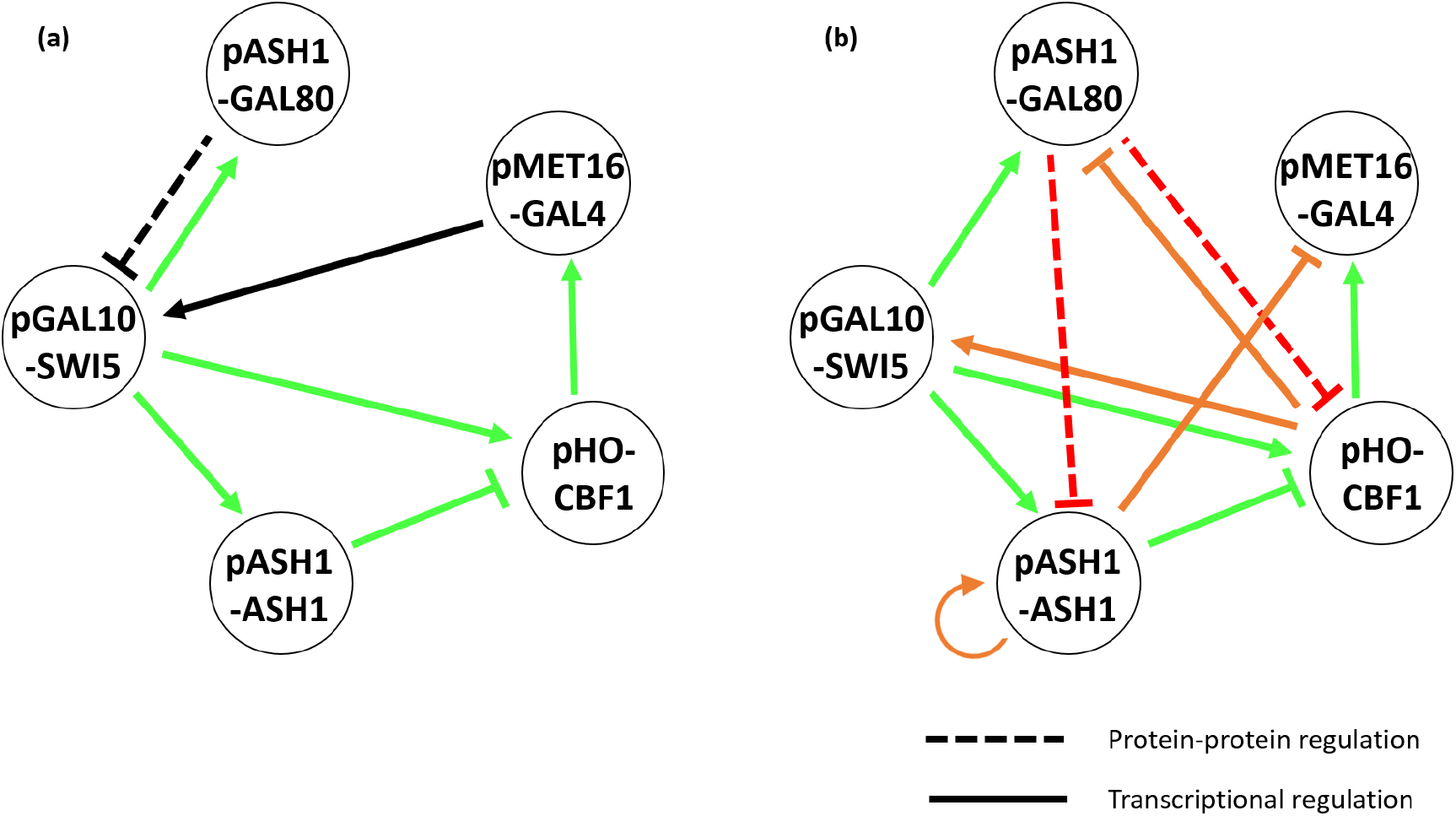
(a) The schematic diagram of the *S. cerevisiae* synthetic circuit. Solid lines represent direct transcriptional regulation and dotted lines indicate indirect transcriptional regulation mediated by a protein-level activation or inhibition of a transcription factor. Edges accurately inferred by our approach are labeled in green, otherwise in black. Gal80 protein can inhibit *SWI5* transcription by preventing Gal4-mediated activation of target genes in the absence of galactose. Modified from the original paper [56]. (b) The schematic diagram of the inferred circuit. Additional inferred edges that are not present in the original design but are supported by previously published literature are labeled in orange. Inferred inhibitory edges indicated in red represent putative mechanisms for indirect repression of *SWI5* by Gal80, but are not supported by known mechanisms of Gal80 function as discussed in the text.

We also tested our model using transcriptional profiles derived from a set of 12 wild-type and targeted TF deletion strains of *Candida albicans*. All strains used in this study are described in Table S3 and are derived from SN156, which is a commonly used derivative of the SC5314 strain that is used widely in *C. albicans* studies [72], [73]. All of the *C. albicans* single TF deletion strains used in this study were reported previously [73]. TF double deletion strains were generated using CRISPR-mediated genome editing to delete the *WOR1* coding sequence as described by Nguyen et al [74]. Steady-state transcript levels were determined using the 3’ quant seq RNA sequencing methodology as described by Moll et al [75]. Briefly, *C. albicans* cells were harvested from mid to late-log cultures and total RNA was isolated using the RiboPure™ RNA Purification kit. cDNA libraries were prepared using the QuantSeq 3’ mRNA-Seq Library Prep kit from Lexogen and multiplexed in pools of 96 libraries. Single-end 100bp reads were obtained using an Illumina HiSeq4000 instrument. The resulting de-multiplexed sequencing reads were trimmed and aligned using STAR Aligner [76] to obtain raw read counts for each transcript genome-wide. The promoter strengths of each gene in the network were determined using capped small RNA (csRNA) sequencing [77]. This method enables the isolation of short nascent mRNA transcripts, rather than full-length mRNAs, and thus provides an instantaneous snapshot of the level of transcriptional activity at each transcriptional start site, genome-wide. Briefly, we enriched for nascent small, capped, RNA molecules from total RNA extracted from mid-log phase *C. albicans* cultures and prepared sequencing ready libraries using the small RNA library preparation kit from New England Biolabs. The resulting libraries were multiplexed and 16 indexed libraries were pooled prior to sequencing on an Illumina HiSeq4000 instrument. Sequencing data were analyzed using HOMER [77]. The mRNA and csRNA sequencing data can be accessed on GEO (GSE217461 and GSE217383). Our evolutionary algorithm, the datasets used in this study, and the results are available on a GitHub repository at https://github.com/UCM-RuihaoLi/GeneRegulatoryNetworkInference. We have also used Zenodo to assign a DOI to the repository: https://doi.org/10.5281/zenodo.7553611.

To account for noise in the experimentally derived transcriptional profiles we measured the “average replicate distance” which describes the average pairwise attractor distance between each of the three biological replicates for each genotypic/phenotypic combination. This metric was then used to determine whether a given GRN model prediction was considered successful, with the basic premise that a successful GRN prediction should yield a transcriptional profile that lies within the average range of noise in the experimentally derived transcriptional profiles. Since some of the experimental replicates had high variance, we also included an attractor distance threshold of 0.16. This threshold was selected based on the performance of a null model, which samples from a uniform distribution with the upper and lower limits as the maximal and minimal expression levels for each gene. For transcriptional profiles with five genes or more, the null model has a 5% chance or less to generate a profile below this cutoff of 0.16 (See Table S4).

## Results

### Consensus GRNs converge upon attractors of reference GRNs

We propose that a successfully inferred GRN should accurately reproduce the experimentally derived attractor states from which it was derived. Therefore, we initiated dynamic simulations using the inferred consensus GRN and the reference GRN at random initial states around their attractor positions. The result showed that their expression levels converged upon the same attractor (Fig. 3b).

### GRN architectures and attractor profiles are strongly coupled

One key hypothesis of our approach is that a GRN’s architecture can be revealed by its attractors. To test this hypothesis under normal circumstances, we arbitrarily generated five *in silico* GRN architectures as test subjects (Fig. 3a) and examined the correlation between their *A_net_* and attractor profiles. Specifically, we randomly mutated the *A_net_* of the reference GRN architectures and observed how their attractor profiles change accordingly. The difference between the mutant GRN architectures 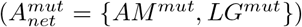 and the reference architectures 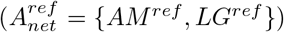 is measured by the Hamming distance, and the difference between their attractor profiles is measured by the attractor distance given by Eq. 10. As shown in Fig. 3c, when the 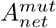 becomes more different from 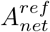, its attractor profiles tend to be further away from the reference. This general trend between *A_net_* and attractor profiles justifies the search strategy of our algorithm, whereby the 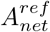 is inferred by gradually mutating *A_net_* and improving the distance between the population’s attractor profiles and those of the reference GRN. In addition, Fig. 3c shows that the attractor distance can reach zero before the Hamming distance goes to zero, indicating that identical attractors can be generated by distinct GRNs. Therefore, even though the reference GRN can be approached by our searching strategy following the trend, it is unlikely to be eventually obtained by an individual architecture in the output, thereby motivating the examination of a consensus GRN.

### Our algorithm correctly infers the GRN architecture in the *in silico* test and outperforms other models

We tested the ability of our model, and several other existing models, to infer GRN architectures using the attractor profiles generated by the five *in silico* reference GRNs depicted in Fig. 3a. To avoid biasing the results in favor of our algorithm, we generated ODEs for these five topologies using SynTReN [57]. We generated simulated transcriptional profiles from the attractors of these ODEs using a global search strategy and utilized them as the input for the *in silico* test. For each instance, we used the attractors as input and ran 600 iterations in algorithm 1. We performed 30 independent and identical inference processes and obtained a consensus *A_net_* by weighted averaging of edge frequencies. We compared the inferred consensus *A_net_* to reference *A_net_* using common machine learning metrics, including the F1 score, area under Receiver Operating Characteristic (ROC) and Precision and Recall (PR) curves. We compared our evolutionary algorithm (EA) method, to six widely used benchmark methods, namely ARACNE [25], CLR [26], GENIE3 [78], MRNET [79], MRNETB [80] and SIMONE [81]). Amongst these methods, only EA and GENIE3 can infer directed networks with asymmetric adjacency matrices, which can differentiate the regulating gene and the target gene (Table S5). Therefore, we symmetrized the inferred networks of EA and GENIE3 as EA_SYM and GENIE3_SYM by making all edges undirected. As presented by Fig. 4, the EA performed generally better than the other algorithms. Furthermore, the protein coordination parameters and basal transcription rate parameters of EA converged well to the ground truth (see supplemental material). None of the examined methods other than EA can infer self-regulatory edges; therefore the diagonal in their inferred adjacency matrix was set to zero. To bridge this difference between methods, we conducted the comparison on an additional set of reference GRNs that contain no self-regulatory edges (Fig. S2). This set of reference GRNs disadvantages our algorithm because other methods are not capable of generating false positives for autoregulation. GENIE3 performed the best out of all methods we considered. Nevertheless, despite the penalty this test imposes on our EA method, it was still among the most competent ones (Fig. S3). Additionally, we observed that when the scale of the *in silico* GRN increases from five to nine genes, it becomes more difficult to infer the *AM*. We believe that additional attractor profiles are needed to reveal additional stable states of large-scale GRNs and to compensate the curse of dimensionality brought by its bigger state space volume.

### Our algorithm can predict novel attractors produced by genetically engineered GRNs

Because the *in silico* inferred and reference GRNs are similar in both structure and associated attractor profiles, we anticipated that attractors predicted by a mutated form of the inferred GRN should closely resemble the attractors produced by an identical mutation in the true GRN. To test this hypothesis, we systematically “deleted” each of the individual genes in the five *in silico* reference GRNs and found their new attractors by searching through the state space. These new attractors are unknown to the inferred GRNs because they had not been used in the GRN inference process. We performed the same gene deletions in the inferred GRNs to see if they could accurately predict the new attractors of the mutated reference GRNs. The attractors of the mutated reference GRNs were generated by SynTReN, while the attractors of the mutated inferred GRNs were predicted by our algorithm. We applied a systematic global search to find attractors and used a random GRN as a control. We found that the single knockout reference GRNs had altogether 384 attractors (combining attractors from all knockouts across all five reference GRNs) and the single knockout inferred GRNs had a combined total of 385 attractors. Of these attractors, 273 (71.1% of the reference GRN attractors and 70.9% of the inferred GRN attractors) were matched. The random GRN showed 32 attractors, none of which were matched. See Fig. 3d for a summary of these results and supplemental material for the complete results.

### Our algorithm revealed unintended edges in an engineered *S. cerevisiae* GRN

To examine how well the GRN dynamic model produced by our algorithm simulates experimentally derived gene expression and to what extent it is robust to measurement noise, we tested our approach using experimental data derived from an engineered synthetic GRN in *S. cerevisiae* [56]. This engineered GRN consists of seven activating or inhibitory edges and five genes, some of which are under control of non-native promoters (Fig. 5 a). Using the experimentally measured promoter strengths and ten distinct steady state gene expression profiles, derived from strains which individually overexpress each of the five genes in galactose and glucose, our model inferred the GRN shown in Fig. 5 b. In glucose, Gal80 blocks Gal4 from activating *SWI5*, while galactose can inactivate Gal80 and Gal4 is free to activate *SWI5*.

Our algorithm correctly identified five of the six transcriptional regulatory edges present in the original design of the engineered GRN (see comparison result in the supplemental material). In addition, our algorithm predicted two additional edges related to protein-protein interactions and four that were not intended in the original design of the engineered GRN, but for which there is experimental evidence in the literature (see Table 1). We believe the missing transcriptional regulation of *SWI5* by Gal4 can be explained as follows. First, as shown in Fig. 3c, the same set of attractors can be produced by different GRNs. In this case, the activation of *SWI5* by a feedback loop via *CBF1* and *GAL4* is replaced by a more direct activation by *CBF1* only. Second, the difference in the regulatory effects of Gal80 is related to how protein-protein interactions are encoded in our ODE framework. Special care must be taken in interpreting protein-protein interactions in the context of the inferred network produced by our algorithm. Our algorithm does not incorporate explicit protein-protein interactions, such as the interaction between Gal80 and Gal4, which leads to the downregulation of *SWI5* and furthermore *ASH1* and *CBF1*. Thus, in our inferred network, the inhibitory edge from *GAL80* to *SWI5* is not present. Instead, this protein inhibition is incorporated into the regulatory function for the targets of Swi5. Specifically, the Swi5 protein activates *CBF1* and *ASH1* transcription, but the protein Gal80 interferes with this activation. Therefore, at the mRNA level, increased *GAL80* transcription does not directly decrease *SWI5* mRNA production; rather it decreases *ASH1* and *CBF1* transcription. Thus, the inhibitory effect of Gal80 on the Swi5 protein is represented as the two inhibitory edges from *GAL80* to *ASH1* and *CBF1*. With this consideration in mind, only the self-activation of *ASH1*, the inhibition of *GAL4* by *ASH1*, the activation of *SWI5* by *CBF1*, and the inhibition of *GAL80* by *CBF1* represent regulatory effects that are not present in the intended synthetic system. We found previous experimental evidence for all these interactions in the literature (see Table 1).

**Table 1:** Experimental evidence for regulatory associations in the synthetic circuit

|  | Cbf1 | Ash1 | Gal4 | Gal80 | Swi5 |
| --- | --- | --- | --- | --- | --- |
| pHO-CBF1 | Inhibition<br>[83] | Inhibition<br>[84, 85] |  |  | Activation<br>[86, 87] |
| pGAL10-SWI5 | Activation<br>[83] | [82] | Activation<br>[88, 89] | Inhibition<br>[90] |  |
| pMET16-GAL4 | Activation<br>[83, 91] | [82] |  |  |  |
| pASH1-GAL80 | Activation<br>[83] | [82] |  |  | Activation<br>[84] |
| pASH1-ASH1 | Activation<br>[83] | [82] |  |  | Activation<br>[84] |

Column names are the TF proteins and row names are the promoters followed by their open reading frames. Orange shaded boxes indicate potential regulatory associations found in YEASTRACT+; blue shaded boxes are experimental evidence found by microarray and/or Northern blot experiments; pink shaded boxes are experimental evidence found by both microarray/Northern blot and ChIP/EMSA experiments.

Furthermore, while investigating the source of these additional edges, we observed that certain elements of the experimentally derived transcriptional profiles did not appear to be consistent with the intended design of the engineered GRN as described by Cantone et al. [56]. Specifically, Cbf1 was intended to serve as the sole activator of *GAL4*, which in turn was meant to serve as the sole activator of *SWI5*. This would imply that, at steady state, *SWI5* should be expressed if and only if Cbf1 is elevated and *GAL4* is expressed. The experimentally derived transcriptional profiles contradict this. They indicated that at steady state, *SWI5* was activated even when *GAL4* was not expressed. Cantone et al. argued that *GAL4* is transiently expressed during an early phase of the experimental protocol, and that the Gal4 protein could persist to activate *SWI5* even after *GAL4* mRNA levels drop. However, this argument contradicts the steady state assumption of the transcriptional data and furthermore does not explain why *GAL4* mRNA levels were low when *CBF1*, which was intended to activate *GAL4*, was overexpressed.

We speculate that these discrepancies between the intended engineered GRN and the experimentally derived data may be explained by unintended regulatory interactions that modify the GRN structure and dynamics. By performing a systematic search on each TF-promoter pair in the intended engineered GRN using YEASTRACT+ [82], we uncovered support for this hypothesis. Specifically, we found experimental evidence from microarray, Northern blot, ChIP, and electrophoretic mobility shift assay (EMSA) experiments, supporting the idea that Cbf1 and Ash1 proteins regulate more than their intended target genes in the circuit. In fact, all four promoters within the circuit can be responsive to Cbf1 and Ash1 (Table 1).

Our inferred GRN predicted additional regulatory interactions beyond those that were intended in the synthetic regulatory network [56], and we identified experimental support for these putative regulatory interactions (Table 1). We conclude that the inferred GRN may have identified actual regulatory associations that impacted the experimentally derived transcriptional profiles, thus allowing our inferred GRN to accurately reproduce the experimentally measured attractor states and resolve the conflict between the intended GRN and the experimentally derived transcriptional profiles. This conclusion is supported by the observation that the attractors reproduced by our inferred GRN have 25.8% of the attractor distance of the mathematical model built by Cantone et al. (supplemental material). Furthermore, the experimental transcriptional profiles showed that *SWI5* was repressed during overexpression of *GAL80* in both galactose and glucose, which was inconsistent with the intended GRN. The attractors produced by our model showed a consistent result: *SWI5* was suppressed when *GAL80* was overexpressed in glucose media, and it was expressed when galactose inactivated Gal80. Our model also explains the low expression of *GAL4* under *CBF1* overexpression: when *CBF1* was overexpressed, *ASH1* was activated by Cbf1 by two feed-forward loops (one via *SWI5* and the other via *GAL80*), and Ash1 in turn inhibited *GAL4*, lowering its expression. Together these results strongly suggest that our evolutionary algorithm approach to model construction can provide significant insight into the structure and regulatory dynamics of “real world” *in vivo* GRNs.

### Modeling the white-opaque switch GRN in *C. albicans*

To expand beyond our model testing using data derived from “known” *in silico* and engineered *in vivo* GRNs, we next applied our algorithm to infer and simulate the dynamics of a naturally occurring GRN which controls reversible differentiation between two distinct cell types—white and opaque—in the human fungal pathogen *C. albicans*. The white and opaque cell types are heritably maintained for hundreds of generations and the frequency of stochastic switching between these two cell types is controlled by a complex, highly interwoven series of transcriptional regulatory interactions [72]. The white and opaque cell types differ in the expression of approximately 18% of all genes in the *C. albicans* genome, thus providing two very distinct attractor states for the underlying GRN. To model the white-opaque GRN, we utilized transcriptional profiles derived from wildtype white and opaque cells, along with a series of strains that lack one or more of the TFs controling the switch (See Table S3). These additional TF deletion strains serve to provide additional steady-state attractors to further constrain the GRN structures. The majority of these strains can switch reversibly between the white and opaque cell types, thus providing two distinct attractor states per strain, with the exception of those TF deletion strains that are “locked” in one cell type or the other. In total, we obtained RNAseq data for seventeen distinct genotypic/phenotypic combinations including two wildtype strains, thirteen single TF deletion strains, and two double TF deletion strains. Each of the deleted TFs is known to impact the frequency of switching between the white and opaque cell types, and is known or predicted to impact the transcriptional profile of the resulting white and/or opaque cell types.

We first tested the ability of our evolutionary algorithm to predict the “unknown” transcriptional profiles produced by the GRNs of the wildtype and single TF deletion strains by omitting the transcriptional profile(s) of a specific genotype from the training dataset and allowing the model to predict the omitted transcriptional profile(s). Transcriptional profiles from the two double TF deletion strains were excluded from all training sets and were reserved as final test subjects for a “fully trained” version of the model developed using the full complement of fifteen wildtype and single TF deletion strain transcriptional profiles as the training dataset. If the attractor distance between the predicted and omitted transcriptional profile(s) was below the average replicate distance, or a cutoff of 0.16, the prediction would be considered successful. The cutoff of 0.16 was selected by the null model, which has less than a 2% chance of generating a result below this cutoff for eight and nine gene networks (See Table S4). Since the null model produces transcriptional profiles by simply picking a value between the maximal and minimal expression levels, while the GRN dynamic system generates transcriptional profiles by numerically solving the differential equations, potential discrepancies may exist between the two. To rule out potential discrepancies due to the GRN dynamic system, we also generated 10,000 random GRNs as a control group and performed the same predictions on the omitted transcriptional profiles. Generally, half of the random GRNs produced fixed-point attractors, while the other half did not reach a steady state. Both the null model and the control GRNs showed similar distribution on their attractor distances and had an average of approximately 0.3.

Overall, nine out of the fifteen omitted wildtype and single TF deletion strain transcriptional profiles were successfully predicted by our model (Table 2). Of these nine successful predictions, eight had an average attractor distance of less than 0.16, and the last one (Δ/Δ*efg1*; Table 2) had an attractor distance above 0.16 but below the average replicate distance, meaning that the predicted transcriptional profiles were within the range of noise in the experimentally derived transcriptional profiles for the *EFG1* deletion strain. The six remaining prediction results showed either attractors exceeding the cutoff, or no attractor at all (indicated by dashes). We note that several of the experimentally derived transcriptional profiles had unusually high variability, as indicated by high average replicate distance values (Δ/Δ*wor3* opaque, Δ/Δ*czf1* opaque, and Δ/Δ*rbf1* opaque; Table 2). This high variability suggests excessive noise in the RNAseq libraries, or multiple states/oscillations existing in these specific cell types, either of which would violate the model assumption of a single stable-state transcriptional profile and make it challenging to evaluate the prediction. If we exclude these highly variable samples, the success rate of the model predictions increases to 66.7%.

**Table 2:**
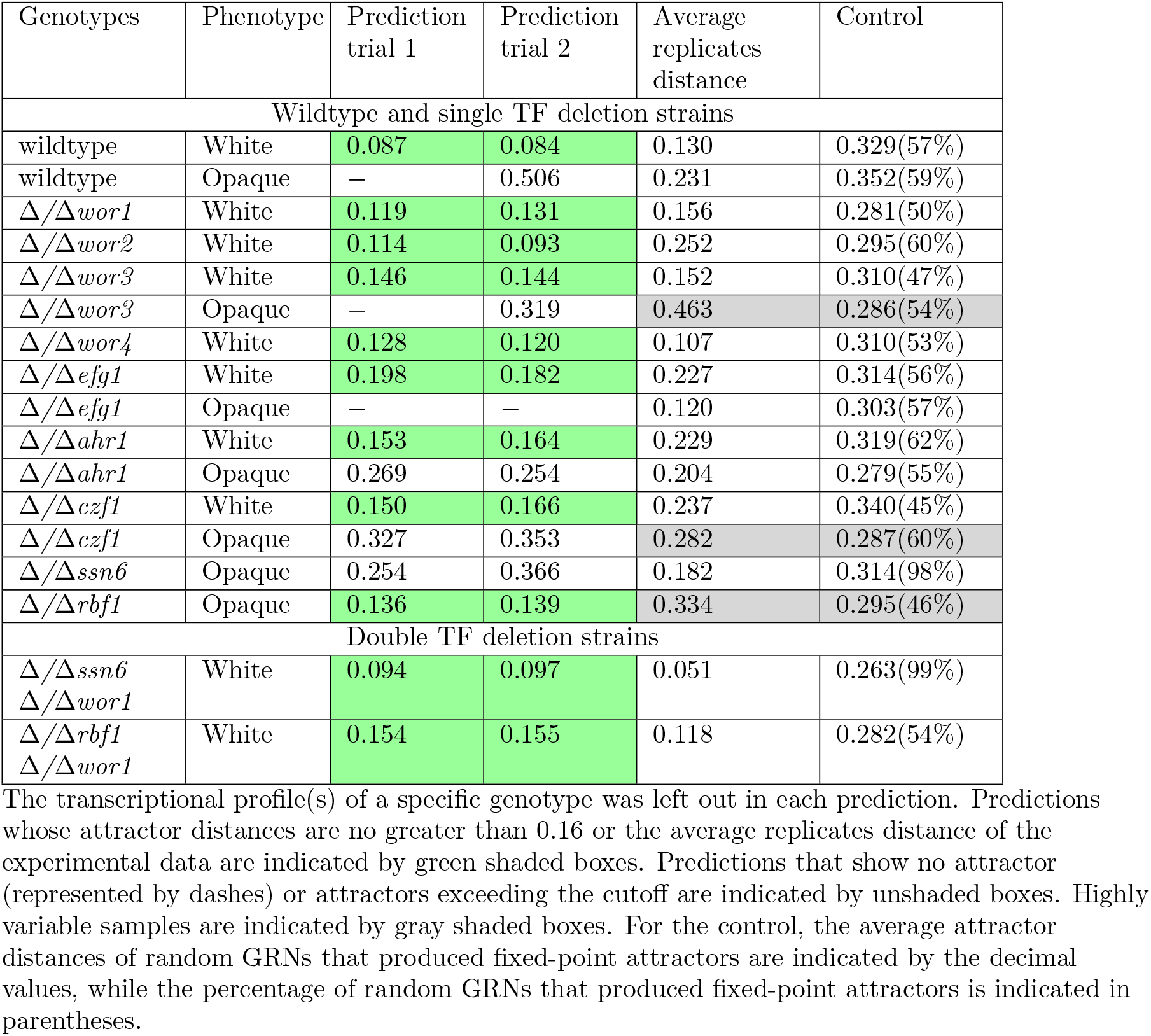
*C. albicans* strains transcriptional profiles prediction results

Next, we applied all fifteen of the wildtype and single TF deletion strain transcriptional profiles as training data to infer a consensus “fully trained” GRN. This consensus fully trained GRN was derived from thirty inferred GRN architectures and then used to predict the transcriptional profiles for two distinct double TF deletion strains. Since more attractors were used in the input, we anticipated that this consensus fully trained GRN should have a higher predictive power than the partially trained model. Both double TF deletion predictions were successful (Table 2), indicating that the transcriptional profiles produced by the consensus fully trained GRN closely mirror the experimentally derived transcriptional profiles for these two strains. Given the predictive accuracy of the consensus fully trained GRN, we next asked whether the underlying architecture, or adjacency matrix, of the inferred GRN also closely resembled the experimentally determined binding interactions between these regulators and their respective coding genes, as previously reported [72, 92, 93]. The GRN architectures inferred by the fully trained model are relatively diverse, with an average success rate of approximately 50% in predicting the experimentally determined TF-gene binding interactions observed in the ChIP data (Fig. S4). This discrepancy is not entirely unexpected given that our *in silico* testing demonstrated that multiple distinct GRN structures, covering a wide range of hamming distances, are capable of producing virtually identical transcriptional profiles, or attractor distances (Fig. 3c).

Given the enormous number of potential GRN architectures in the search space, and the fact that distinct GRNs, which produce identical attractors cannot be differentiated based purely on transcriptional profiles, we asked whether incorporating TF binding constraints could enable the model to converge upon an architecture that more closely resembles the experimental ChIP data while simultaneously reproducing accurate transcriptional profiles. To bias the model toward the GRN architecture observed in the experimental data, we included a TF binding probability function in our evolutionary algorithm. Briefly, this function alters the probability of an edge being created or removed in the adjacency matrix, thus biasing the inferred GRNs towards the experimentally determined architecture. However, if the resulting GRNs fail to converge upon the experimental attractors, the evolutionary algorithm would ultimately converge upon a distinct GRN structure if needed to fit the transcriptional profiling data. We applied all seventeen of the transcriptional profiles used above, plus the previously published *in vivo* TF-DNA binding data, to infer “directed” GRNs. On average, the individual directed GRNs retained approximately 90% of the experimentally determined TF binding interactions while also reproducing most of the experimentally derived transcriptional profiles (Fig. S4). The consensus directed GRN, constructed by the high-frequency edges of the individual directed GRNs, accurately reproduced thirteen out of the seventeen experimentally observed transcriptional profiles and eighty out of the eighty-one physical binding interactions between each of the regulatory TFs and their respective coding genes. The transcriptional profiles that the consensus directed GRN failed to incorporate were wildtype opaque, *Δ/Δwor3* opaque, *Δ/Δahr1* opaque, and *Δ/Δssn6* opaque, most of which had relatively high variability in their biological replicates (see full report for both *in silico* and *in vivo* prediction tests and inferred GRNs in the supplemental material). Together these results indicate that it is indeed possible to converge upon a GRN structure that closely mirrors the experimentally determined TF-DNA binding data for the white-opaque switch, while accurately producing many of the same attractor states observed *via* RNAseq. However, this data also suggests a high degree of redundancy or potential for plasticity within the white-opaque GRN, thus compromising the ability of our model to infer the observed GRN structure based solely on transcriptional profiling data.

## Discussion

In this work, we extended the attractor-matching strategy from a Boolean model to an ODE-based model by incorporating transcriptional kinetic parameters. We consider transcriptional profiles of stable cell types as fixed-point attractors [94] in the mRNA state space, and search for the GRN architecture that produces these attractors. We found in the *in silico* simulation that GRN architectures are significantly correlated with the attractors they produce. This correlation supports the logic of applying the attractor-matching approach to GRN inference. The ability of our approach to infer “unknown” GRNs has been validated using both simulated datasets derived from “known” *in silico* GRNs and *in vivo* test datasets from an engineered GRN in *S. cerevisiae*. Our approach outperformed six other leading GRN inference methods when applied to the *in silico* attractors generated by SynTReN ODEs. In the *in vivo* test, our approach not only successfully identified five of the six intended transcriptional edges, but also revealed some unintended edges that might account for the inconsistency between the designed GRN and experimentally derived transcriptional profiles. In addition to inferring GRN architecture based on transcriptional profiles, our approach can also predict the effects of genetic perturbation on the inferred GRN. As a proof of principle, we used the inferred GRNs generated during our *in silico* model testing to then predict the unknown attractors that would be produced upon genetic perturbation of the original reference GRN (i.e., by deletion of each TF). The inferred *in silico* GRNs successfully predicted 71.1% of the attractors produced by the reference GRNs using the identical knockout strains (Fig. 3d), indicating that our approach can effectively capture GRN behavior based on transcriptional profiles. This result further suggests that our approach can be used to generate testable predictions on the behavior of *in vivo* GRNs. Specifically, we envision the application of this approach to a hybrid computational and *in vivo* experimental process whereby GRNs are inferred based on *in vivo* transcriptional profiles, the inferred GRNs are perturbed *in silico* to generate “mutant” transcriptional profiles, and the accuracy of inferred GRNs are ultimately assessed by comparing predicted versus observed transcriptional profiles generated using *in silico* versus *in vivo* mutant strains. The accuracy of the inferred GRN could thus be supported if the predicted and experimentally measured transcriptional profiles converge. If not, the *in vivo* mutant strain and the resulting experimentally derived attractors could reveal a new pattern of GRN dynamics that had not been covered by the initial input attractors, and would thus complement the original wildtype attractors to further refine the inferred GRN. In this manner, it should be possible to iteratively refine predicted GRNs until they approximate the *in vivo* results.

As a proof of principle, we applied this iterative computational and experimental strategy to infer the GRN governing the white-opaque switch in *C. albicans*. We first used a dropout strategy to infer GRNs based on a subset of available data and tested the ability of the inferred GRNs to predict the transcriptional profiles that were omitted from the training data. This approach led to an overall success rate of 66.7%, which approaches the 71.1% success rate observed in our *in silico* testing. Next, we demonstrated that a “fully trained” GRN inferred from all fifteen of the wildtype and single gene deletion strain profiles was successful in predicting the transcriptional profiles of two distinct “unknown” double TF knockout strains that were omitted from our training data. This result demonstrates that the inference of a GRN using a set of known attractors can bring insight into attractors that exist biologically but have not yet been measured in the lab. Although the inferred white-opaque GRNs accurately predicted most if not all of the dropped-out transcriptional profiles, they did not fully converge upon the TF localization patterns that we have observed in *in vivo* genome-wide TF localization experiments. To further constrain the white-opaque GRN, we inferred the “directed” GRNs with all seventeen available experimentally measured transcriptional profiles and included ChIP data that biases the GRN architecture towards the TF localization pattern observed *in vivo*. This consensus directed GRN accurately reproduced 76% of the RNAseq-derived transcriptional profiles and converged upon 99% of the ChIP-derived TF binding interactions. Although this directed GRN model focuses on the small-scale regulatory network of transcription factors that underlie the white-opaque switch, it should be feasible to expand the size of the network by considering the transcription factors as hub genes, which are highly correlated with other non-TF genes in a biologically significant module. Typically, the expression patterns of non-TF genes can be expressed by a linear combination of transcription factor expression patterns [95]. With the TF regulatory network accurately inferred, it should be possible to classify the interactions amongst biological modules and include the regulation of downstream target genes in the network model. This hybrid regulatory network model could be iteratively tested and refined to provide further insights into the transcriptional regulatory dynamics of this highly intertwined and complex GRN that controls cellular differentiation in *C. albicans*.

There are potential pitfalls that can impact our approach. First, regulatory elements other than TFs, such as non-coding RNA molecules, post-translational modifications, and chromatin modifiers/remodelers, can also influence the behavior of a GRN of interest. Their regulatory effects can lead to false compensatory TF regulations and make the inferred network converge less often. Second, as observed in our *C. albicans* GRN modeling, noise in the experimental data can lead to a “fuzzy” target for prediction and compromise the ability of the approach to fit the transcriptional profiles into a GRN. Furthermore, while RNAseq data derived from a particular genotypic/phenotypic state is assumed to represent a fixed-point attractor, this is not necessarily the case. Multiple stable states or oscillatory transcriptional outputs could exist within a population of cells that appear to be phenotypically homogeneous, thus bulk RNA sequencing could average out single-cell heterogeneity and underlying GRN dynamics. These limitations could lead to inferred GRNs that simulate biased or non-existent targets. Third, our approach assumes that the highest expression levels for each gene have been observed in the input transcriptional profiles and utilizes them to estimate the unknown parameters. Potential bias in parameter estimation can occur if this assumption is not satisfied. Moreover, our approach has simplified the way that multiple activators or inhibitors regulate a target gene, either independently as monomers or cooperatively as a polymer, but *in vivo* TFs could have more complex and sophisticated forms of incorporation than modeled in our approach. These and likely other confounding factors have the potential to adversely impact the process of GRN inference and can cause reduced accuracy in predicting unknown transcriptional profiles.

The most significant challenge in GRN inference is perhaps the inherent functional redundancy and plasticity of real-world GRNs. This was apparent in our *in silico* testing where we observed that GRNs differing in as many as ten regulatory interactions can produce qualitatively similar transcriptional profiles (Fig. 3c). Similarly, we observed that most of the attractors produced by the *C. albicans* white-opaque GRN could be reproduced, and “unknown” attractors predicted, even when the inferred GRN does not closely match the experimentally determined GRN architecture (Fig. S4). These results are consistent with the idea that GRN structures can evolve while maintaining the same overall output, which is also supported by experimental evidence [96]. For example, Tsong et al. [97] identified a set of sexual differentiation genes that are negatively regulated in *S. cerevisiae*, but are believed to have been positively regulated in an ancestral fungal species. In this example, the overall output of the transcriptional circuit remains the same, despite significant changes in GRN architecture. Our work provides a mathematical foundation for the idea that GRN architecture has plasticity and evolves [98–100] under selective pressure [101]. Thus, experiments performed under a specific set of experimental conditions may fail to reveal some of the evolutionary pressures that have constrained the behavior of real-world GRNs under distinct environmental conditions. While the impact of these unobserved evolutionary pressures on GRN architecture and logic could be revealed by extensive measurements of GRN output in an array of different environmental conditions, we hypothesize that the iterative model refinement strategy that we propose here may represent an efficient alternative strategy.

Despite these potential limitations and challenges, we have shown that our approach outperforms competing GRN inference tools when applied to *in silico* datasets and has a unique set of capabilities that can provide insights into the inner workings of *in vivo* GRNs. In future iterations of our GRN inference approach, we can incorporate other types of interactions between TFs that are not independent as assumed by default. For instance, to consider the fact that Gal80 can only perform gene regulation by binding to Gal4, we can add the following rule to the algorithm: if a target gene is regulated by Gal80 but not by Gal4, the regulation of Gal80 on this gene will be voided. Interactions between metabolites and TFs, such as IPTG deactivating the lac repressor, can also be incorporated into the approach by adding similar rules. In this manner, our approach can flexibly integrate more detailed biological information beyond sequencing data and better simulate complex biological systems. We have demonstrated that our GRN inference approach can successfully narrow down the number of potential GRN structures for a given transcriptional program using only a relatively small number of transcriptional profiles. For example, there are 3^81^ potential adjacency matrices and 2^18^ protein coordination matrices for a GRN with nine genes in our framework, not to mention the number of possible values of *f*_0_s bounded between 0 and 1. In total this creates a GRN search space in excess of 1.2 × 10^44^ distinct configuration options. In the *in silico* test of a nine-gene network, the GRN inferred by our approach was at the 5.6 × 10^−13*th*^ percentile in a network quantity distribution based on the Hamming distance to the reference GRN, whereby most of the incorrect networks were eliminated. Therefore, our approach can effectively select candidate GRNs that can be further examined by experimentation. Based on these results, we believe that this model is a valuable companion to experimental approaches for deciphering the structure and logic of *in vivo* GRNs.

## Acknowledgements

We gratefully acknowledge computing time on the Multi-Environment Computer for Exploration and Discovery (MERCED) cluster at the University of California, Merced. We especially extend our appreciation to Dr. David Ardell who provided valuable advice on the development of our approach. We also thank all Hernday and Nobile lab members for providing insight on the project.

## Author contributions

A.D.H. and R.L. conceived the project with early input from S.S.S. A.D.H. managed and supervised the project. R.L. formulated, analyzed, and interpreted the GRN inference model with key input from J.C.R. and additional advice from A.D.H., R.A., C.J.N., and S.S.S. *C. albicans* RNA sequencing data was generated by M.M.Q. and M.N.Q. and R.L. performed data analysis with input from A.D.H., M.N.Q. and M.M.Q. R.L. wrote the first draft of the manuscript and A.D.H. extensively revised the manuscript with input from J.C.R., R.A., and C.J.N. All authors contributed to editing the manuscript.

